# Network Topology reveals disrupted AMPA receptor stabilization leads to an impaired LTD in epileptic synapses

**DOI:** 10.64898/2026.08.10.743452

**Authors:** Pacheriyil Meghna, Suneel Kateriya, Pradeep Punnakkal

## Abstract

Cognitive comorbidities in epilepsy patients may be the result of synaptic alterations and impaired synaptic signalling. Electrophysiological evidence demonstrates that epileptic synapses undergo a GluN2B-dependent metaplastic shift, where a low-frequency stimulation protocol unexpectedly induced long-term potentiation (LTP) rather than long-term depression (LTD). However, the downstream postsynaptic structural cascade responsible for this functional impairment remains unresolved. To elucidate the molecular architecture driving this shift, this study employed an *in silico* protein-protein interaction network approach using Cystoscape. A baseline intersection network of LTD and epilepsy-associated genes were constructed, anchored with GRIN2B, and topologically ranked to identify hub proteins. This analysis identified a core module biased toward synaptic potentiation, dominated by the kinase CAMK2A, AMPA receptor subunits, and auxiliary Transmembrane AMPA Receptor Regulatory Proteins (TARPs) and CNIH2. These provided a structural basis for the prolonged receptor retention and delayed deactivation kinetics characteristic of epileptic synapses. Mapping the LTD-execution machinery against this interactome revealed that calcineurin was topologically segregated and lacks direct connectivity from the central AMPA receptor complex. Further studies would be required to test and confirm the involvement of these proteins. To experimentally validate these *in silico* findings, human transcriptomic data from cortical and hippocampal tissues of drug-resistant epilepsy patients was also analyzed which confirmed the significant upregulation of CNIH2 and CACNG2 in both tissue types. The cross-validation with patient transcriptomic data, demonstrated that the epileptic synapse undergoes a pathological shift. Hence, the upregulation of the auxiliary proteins functionally overpowers the established LTD machinery and prevents LTD consolidation.

## 1. Introduction

Epilepsy is one of the most common chronic brain conditions characterised by recurrent, unprovoked seizures due to an imbalance in the excitatory and the inhibitory neurotransmission leading to an abnormal and hyper-synchronous neuronal activity in the brain ^1^. It affects over 70 million people of all age groups across the globe, increases the risk of premature death up to three times, and imposes a major economic, psychosocial, and physical burden on families and health systems ^2^.

Learning and memory deficits and cognitive impairments cause a significant burden to the Patients with Epilepsy (PWE). Long-term potentiation (LTP) and long-term depression (LTD) are the extensively studied cellular mechanisms of memory formation encoded by changes in the synaptic plasticity. They are activity-dependent, bidirectional modifications that regulate the synaptic strength. In the hippocampus, the hyperexcitation caused due to uncontrolled seizures leads to cell death, resulting in mossy fibre sprouting, thus resulting in altered synaptic transmission ^3,4^. LTP has been extensively studied and was impaired in animal models of chronic epilepsy ^5,6^. Studies have shown that the standard high-frequency stimulation (HFS) or the theta-burst stimulation (TBS) protocols that induce robust LTP in healthy brain slices fail to elicit further synaptic strengthening in epileptic hippocampal slices ^7^. At the molecular level, this impairment of LTP is driven by excessive calcium influx that keeps kinases like Calcium/calmodulin-dependent protein kinase II (CaMKII) highly active, leading to rapid reinsertion and stabilization of AMPA receptors at the synapse. Therefore, the failure to induce LTD results in a metaplastic alteration, ultimately impairing bidirectional plasticity ^8,9^.

In the epileptiform induced slices, it has been documented that a failure of LTD consolidation while using low-frequency stimulus (LFS) protocol. Instead of inducing LTD in the epileptiform induced brain slices, the plasticity was shifted towards LTP ^9^. The molecular gatekeeper of this shift in the plasticity was identified to be the GluN2B subunit of the NMDA receptors. A previous study confirms the role of this subunit by blocking GluN2B using the subunit-specific antagonist, Ro 25-6981 which inhibited the induction of LTP when given LFS protocol ^9^.

While the trigger for this plasticity shift is clearly GluN2B dependent, the downstream postsynaptic trafficking involving calcium signalling, kinases, phosphatases, and AMPA receptor trafficking, that executes this impairment remains unresolved. To bridge this gap between electrophysiological phenotype and the underlying downstream molecular processes this study utilises an *in silico* connectome approach using STRING (version 12.0) and Cytoscape 3.10.4. This study maps the genes associated with both LTD and epilepsy, and their interaction with GRIN2B, protein encoding GluN2B to get insights into how this metaplasticity could be occurring at the systems level.

## 2. Methods

### 2.1 Identification of Target Proteins and Baseline Network Construction

To map the molecular intersection between pathological network hyperexcitability and altered synaptic plasticity, target proteins associated specifically with LTD (<u>hsa04730</u>) and Epilepsy (<u>DOID:1826</u>) were retrieved from online database STRING (https://string-db.org/) ^10^. The individual networks were then merged using Cytoscape 3.10.4 to get baseline Protein-Protein Interaction (PPI) network ^11^. The species was restricted to *Homo sapiens*, and a minimum interaction confidence score of ≥0.4 (medium confidence) was applied. This medium threshold was chosen intentionally to capture weak but essential signalling events such as kinase activity that drive synaptic plasticity and are often missed by higher cut-offs. To prevent false positives from this medium setting, the network was strictly limited to highly reliable data sources: experimental data, curated databases, and co-expression metrics excluding less reliable sources, such as text-mining and predictive data.

### 2.2 Network Expansion and Topological Hub Identification

As concurrent electrophysiological recordings demonstrated that pharmacological blockade of the GluN2B subunit halted the LTD-to-LTP metaplasticity shift, the GRIN2B protein was introduced into the expanded network of the three hub proteins identified in the LTD/Epilepsy network. The expanded network of the three hub proteins was used here because this study takes into consideration that this metaplastic shift could be due to an interplay amongst the downstream signalling and not just the primary protein set.

After identifying the hub proteins in this network, the CytoHubba plugin was employed so as to systematically identify the most critical regulatory protein within this common protein-set of the 3 core hub protein vs GRIN2B associated proteins. CytoHubba utilizes various topological algorithms to rank nodes by network essentiality. In this study, nodes were evaluated using the Maximal Clique Centrality (MCC) scoring method. MCC was specifically selected because it is mathematically designed to identify and extract the most densely connected, complete topological clusters (cliques) within complex PPI networks, which often correspond to core functional ^12,13^. To define the boundaries of this core module, nodes were ranked by their MCC scores, and the top 8 hubs were selected based on a distinct inflection point in the scoring distribution. This ensured that only the most strictly integrated functional clique was retained for downstream analysis.

### 2.3 Functional Enrichment Analysis

To translate the structural proteomic network data into functional biological mechanisms, Gene Ontology (GO) and Kyoto Encyclopedia of Genes and Genomes (KEGG) pathway enrichment analyses were conducted on the identified hub proteins using the STRING app within Cytoscape, and the resulting data were visualized using GraphPad Prism. GO analysis categorized the proteins by Biological Processes (BP), Cellular Components (CC), and Molecular Functions (MF) ^12^. Enrichment significance was determined with a threshold set at *P* < 0.05. This step was critical for verifying whether the structural hub networks structurally localized to the postsynaptic density and functionally governed AMPA receptor trafficking.

### 2.4 Targeted Sub-Network Construction and Kinase-Phosphatase integration

Guided by literature, a highly targeted sub-network was isolated and constructed in Cytoscape. This targeted connectome analysis was used to map the probable proteins involved in the postsynaptic mechanisms governing this metaplasticity.

To further evaluate the mechanistic balance of plasticity execution within the identified epileptic network, a comparative topological integration was performed. During synaptic plasticity, LTP is driven by the kinase CaMKII, whereas LTD relies on the phosphatase calcineurin. Because our expanded topological analysis unexpectedly identified CAMK2A (CaMKII) as a dominant, highly connected core hub within the GRIN2B-anchored network, it was biologically necessary to assess the opposing LTD machinery.

Therefore, the calcineurin catalytic subunits, Protein Phosphatase 3 Catalytic Subunit α (PPP3CA) and Protein Phosphatase 3 Catalytic Subunit β PPP3CB, representing the standard LTD execution machinery were deliberately mapped against the 8-hub network. This mapping was conducted specifically to determine whether LTD-associated phosphatase signalling could structurally integrate into the core AMPA receptor complex to counterbalance the LTP-associated kinase.

### 2.5 Construction of the Comprehensive Mechanistic Network

To visualize the complete proposed metaplasticity architecture, a final, comprehensive PPI network was constructed integrating all critical proteins identified across the stepwise analyses. This unified interactome consists of the initial baseline intersection hubs, the GRIN2B-anchored expanded core, the targeted network, and the LTD-associated phosphatase machinery (PPP3CA, PPP3CB).

### 2.6 Transcriptomic Validation using NCBI GEO Dataset

To evaluate the clinical and transcriptomic relevance of the identified metaplasticity hubs, human bulk RNA-sequencing data was retrieved from the NCBI Gene Expression Omnibus (GEO) database under the accession number GSE256068. Differential gene expression analysis was conducted using the NCBI GEO2R interactive web tool. Data normalization and statistical analysis were performed using the *limma* R package integrated within GEO2R, applying log2 transformation and quantile normalization. To strictly control for regional transcriptomic variation, samples were physically region-matched. The analysis independently compared epileptic cortical tissue (n=43) versus control cortex (n=12), and epileptic hippocampal tissue (n=63) versus control hippocampus (n=7). According to the overall dataset metadata, the available cortical cohorts exhibited standard sex distributions (Epilepsy pool: 35M/36F; Control pool: 5M/9F) and age demographics (Epilepsy mean age: 11.8 years; Control: 20.6 years). Similarly, the hippocampal cohorts were evaluated for sex (Epilepsy pool: 40M/24F; Control pool: 8M/5F) and age (Epilepsy mean age: 34.7 years; Control: 58.2 years). While regional matching was applied, a limitation of this secondary analysis is that comprehensive clinical metadata regarding post-mortem interval (PMI) and prior anti-seizure medication history was not comprehensively provided in the public GSE256068 repository, preventing statistical adjustment for these specific clinical variables. To validate the proposed network, the expression levels of the core 8 hub proteins, the auxiliary protein CNIH2, and their first-degree interacting proteins were evaluated.

## 3. Results

### 3.1 Identification of Core LTD and Epilepsy Intersection Hubs

On mapping the baseline Protein-Protein Interaction PPI network constructed using the proteins associated with LTD and epilepsy, three hub proteins were identified to be involved in both the physiological processes. These are: GRIA2, PLCB1, and CACNA1A (Fig 1a, and 1b). GRIA2 encodes the AMPA glutamate receptor subunit, GluA2, that dictates the calcium permeability ^13^; PLCB1 encodes the PLC-beta 1 enzyme which breaks down PIP2 in the calcium-dependent signal transduction ^14^; and CACNA1A gene encodes the alpha-1A subunit of calcium channel and is involved in the pore forming of CaV2.1 or P/Q-type voltage-gated calcium channel ^15^.

**Fig 1:**
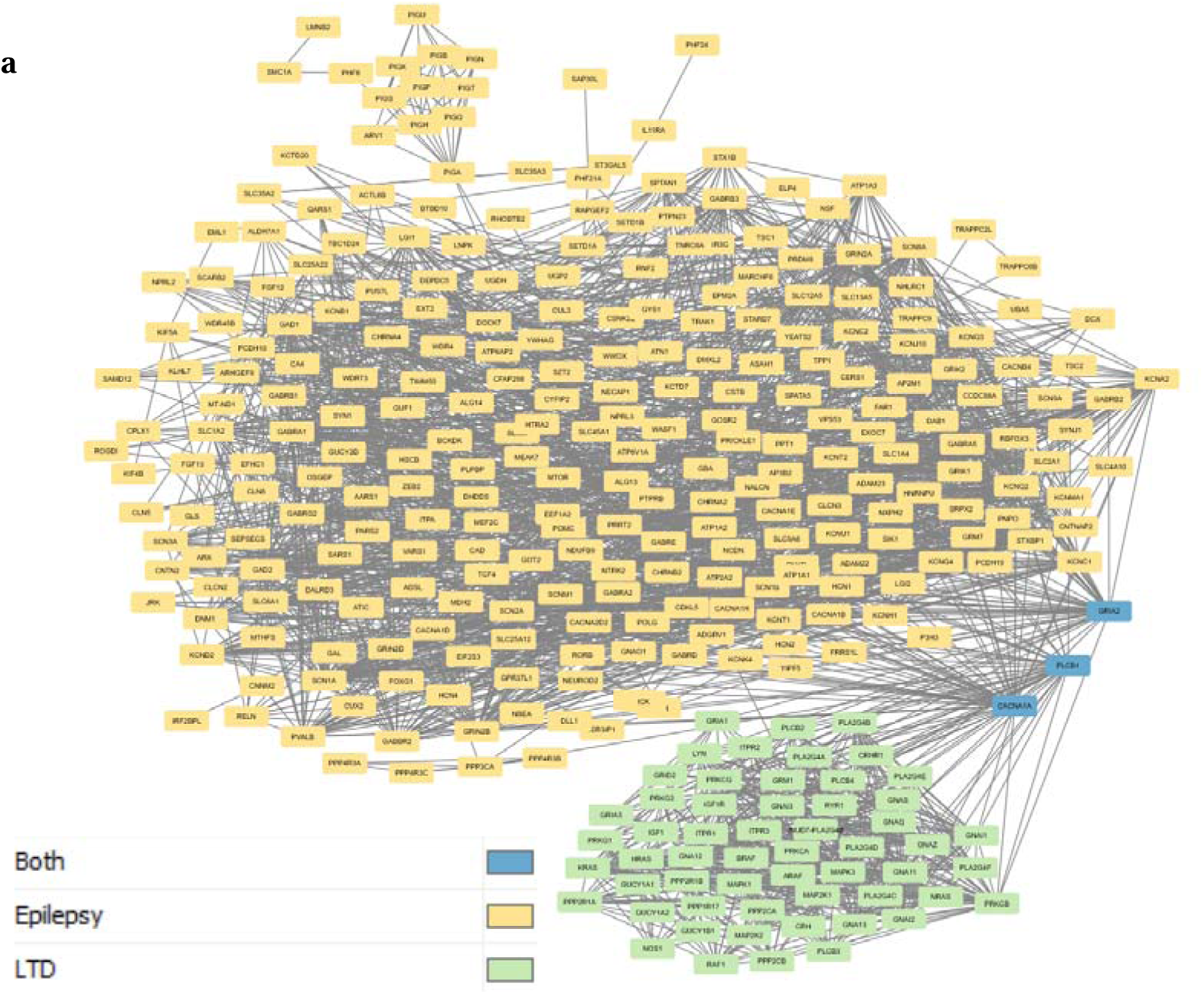
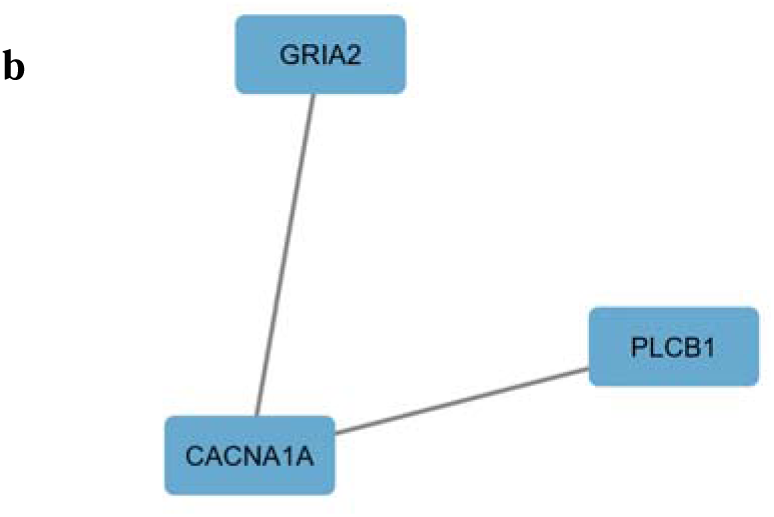
PPI network of proteins involved in epilepsy and LTD constructed using Cytoscape. (a) The green colour represents the proteins of LTD and the yellow colour represents the proteins of Epilepsy. The blue colour shows the common or the hub proteins. This network has 328 nodes and 2243 edges. (b) Extracted network of the hub proteins.

To validate the clinical and physiological relevance of these hub proteins, functional enrichment analyses were performed using Cytoscape and plotted using GraphPad Prism. Disease-gene associations (DISEASES) and Human Phenotype (Monarch) enrichment confirmed links to pathological hyperexcitability, with significant localization to “Epilepsy,” “Focal-onset seizure,” and “Motor seizure” phenotypes (Fig 2a and 2b).

**Fig 2:**
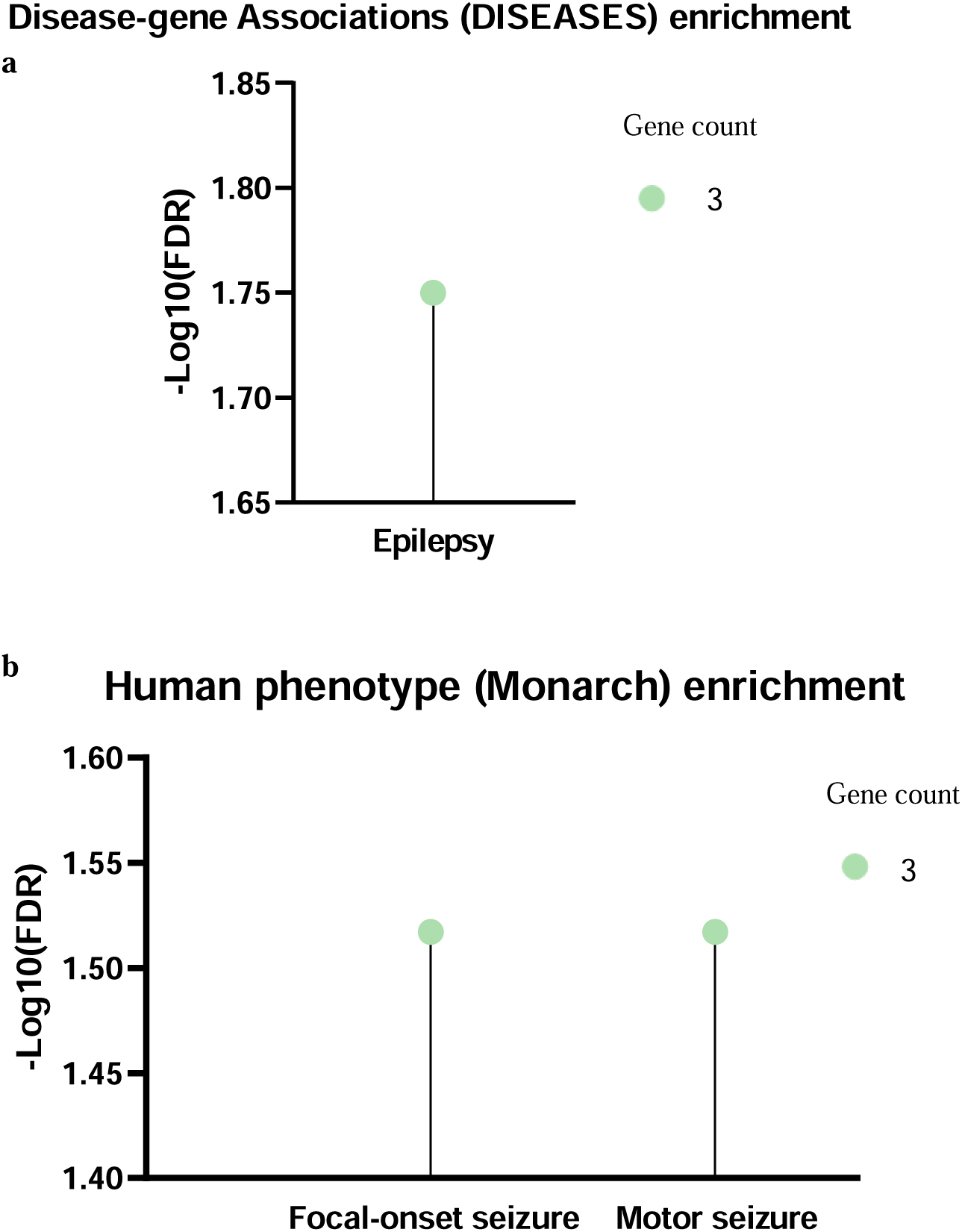
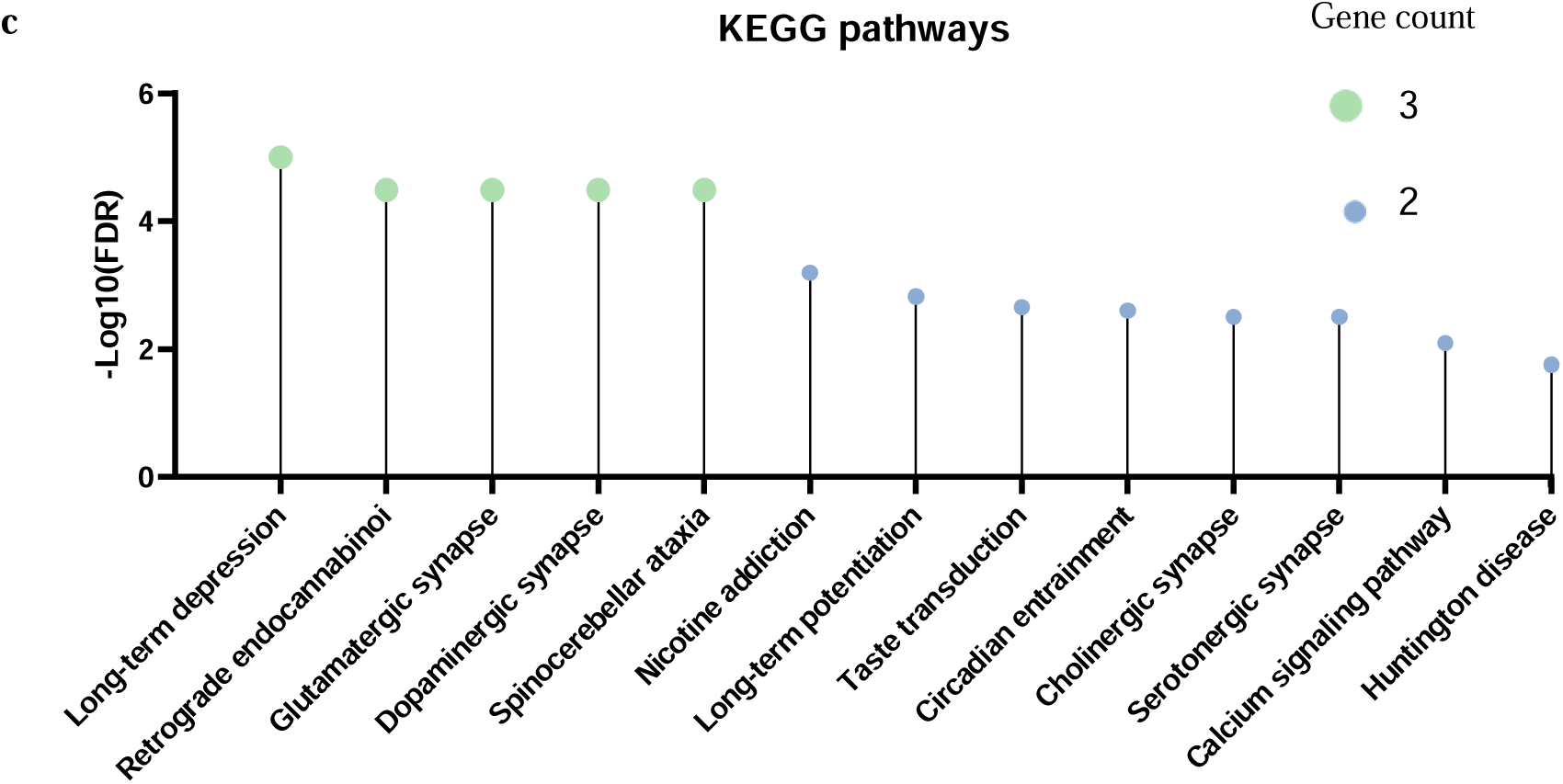
Functional enrichment analysis of the three hub proteins extracted via Cytoscape and visualized using GraphPad Prism. (a) Disease-gene Associations (DISEASES): The network shows highly significant enrichment for general epilepsy. (b) Human Phenotype (Monarch) Enrichment: The hub proteins are significantly associated with specific clinical manifestations of network hyperexcitability, notably focal-onset and motor seizures. (c) KEGG Pathway Enrichment: Analysis confirms the biological localization of these proteins to the glutamatergic synapse and long-term depression.

Simultaneously, KEGG pathway enrichment demonstrated that these core proteins predominantly govern “Long-term depression” and general “Glutamatergic synapse” activity (Fig 2c). Although the network also showed enrichment for ‘Long-term potentiation’, this could be attributed to the shared upstream calcium channel and basic AMPA receptor configuration roles. Hence, the isolation of GRIA2, PLCB1, and CACNA1A established a validated, calcium- and AMPA-dependent molecular foundation bridging LTD and epilepsy.

### 3.2 GRIN2B-Anchored Network Expansion Reveals a Metaplasticity Bias Toward AMPAR Stabilization

Taking the electrophysiological findings into consideration, the GRIN2B associated protein network was then introduced into the expanded network of the three hub genes (Fig 3a). By applying the MCC algorithm, the most highly integrated topological network within this expanded network was successfully extracted. Based on the highest MCC scores prior to the scoring drop-off, a core module of eight interacting proteins was isolated (Fig 3b and 3c). As expected from the MCC methodology, this generated a densely connected sub-network; however, the critical biological finding was the identity of these core hubs, which were heavily dominated by Transmembrane AMPA Receptor Regulatory Proteins (TARPs) namely CACNG8, CACNG3, CACNG2, CACNG5, and CACNG4, and AMPA receptor subunits GRIA4 and GRIA2, alongside the central kinase CAMK2A.

**Fig 3:**
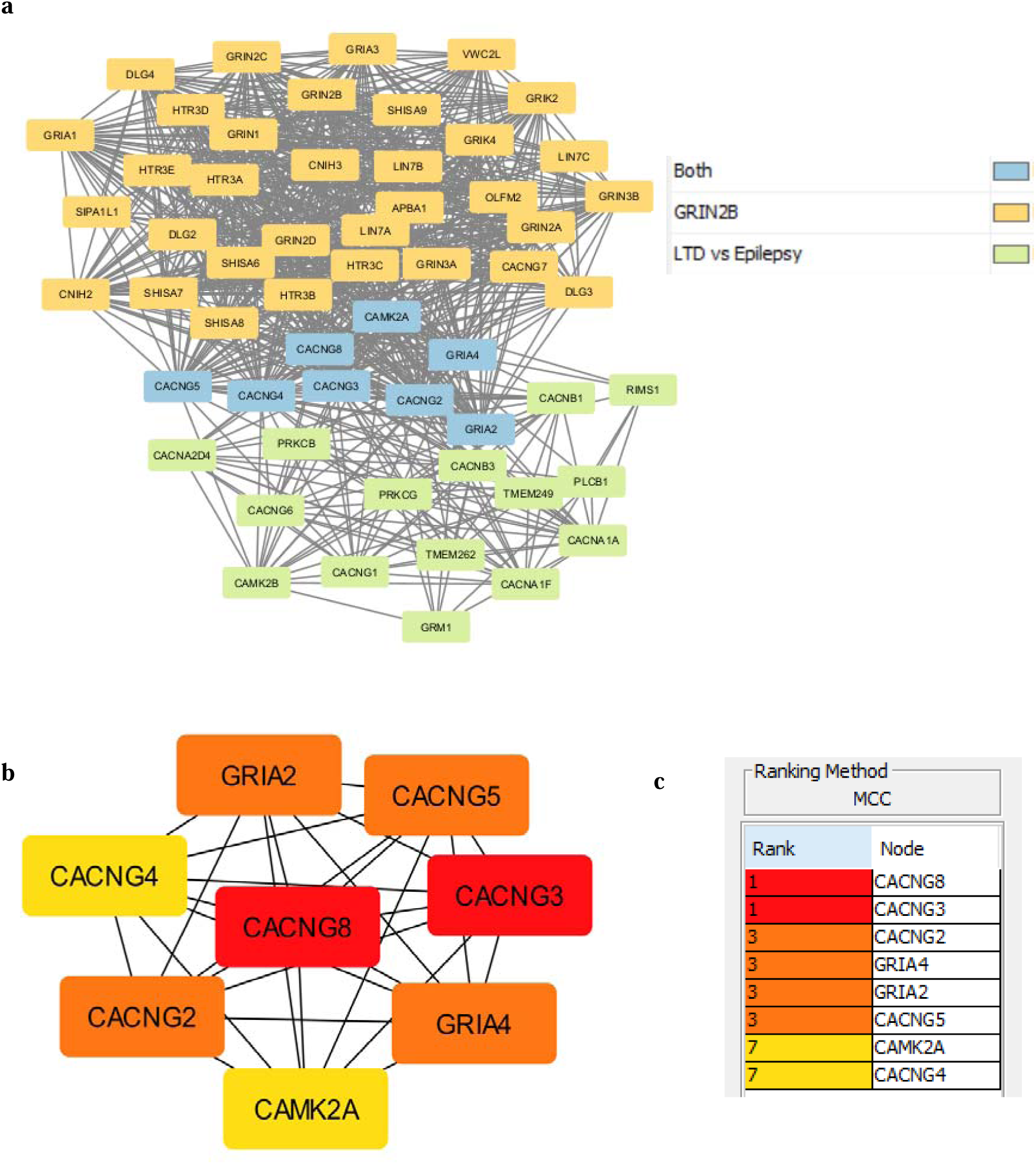
GRIN2B-anchored network expansion and identification of the AMPA receptor-stabilizing core module. (a) A comprehensive protein-protein interaction (PPI) network constructed by expanding the baseline LTD–epilepsy intersection hubs and using GRIN2B as a seed node to model NMDA-dependent plasticity signalling. Nodes are color-coded by their network origin (yellow: GRIN2B-associated; green: LTD and epilepsy baseline hubs; blue: shared intersecting nodes). This network has 56 nodes and 764 edges. (b) The core sub-network comprising the top eight essential hub proteins extracted from the expanded interactome having 8 nodes and 28 edges. (c) Topological ranking of the network nodes by MCC scoring algorithm.

To evaluate the structural capacity of this core module to support plasticity shifts, functional enrichment analysis was conducted. Reactome pathway analysis of the 8 hub proteins revealed a high confidence involvement of these proteins in the ‘Trafficking of AMPA receptors’ and ‘Activation of AMPA receptors’, alongside ‘Long-term potentiation’. Although the ‘Trafficking of the AMPA receptors’ could indicate a receptor internalisation, the presence of ‘LGI-ADAM interaction’ indicates the trafficking of AMPARs towards the post-synaptic membrane via PSD-95-mediated increase of AMPARs at the post-synaptic membrane (Fig 4a).

**Fig 4:**
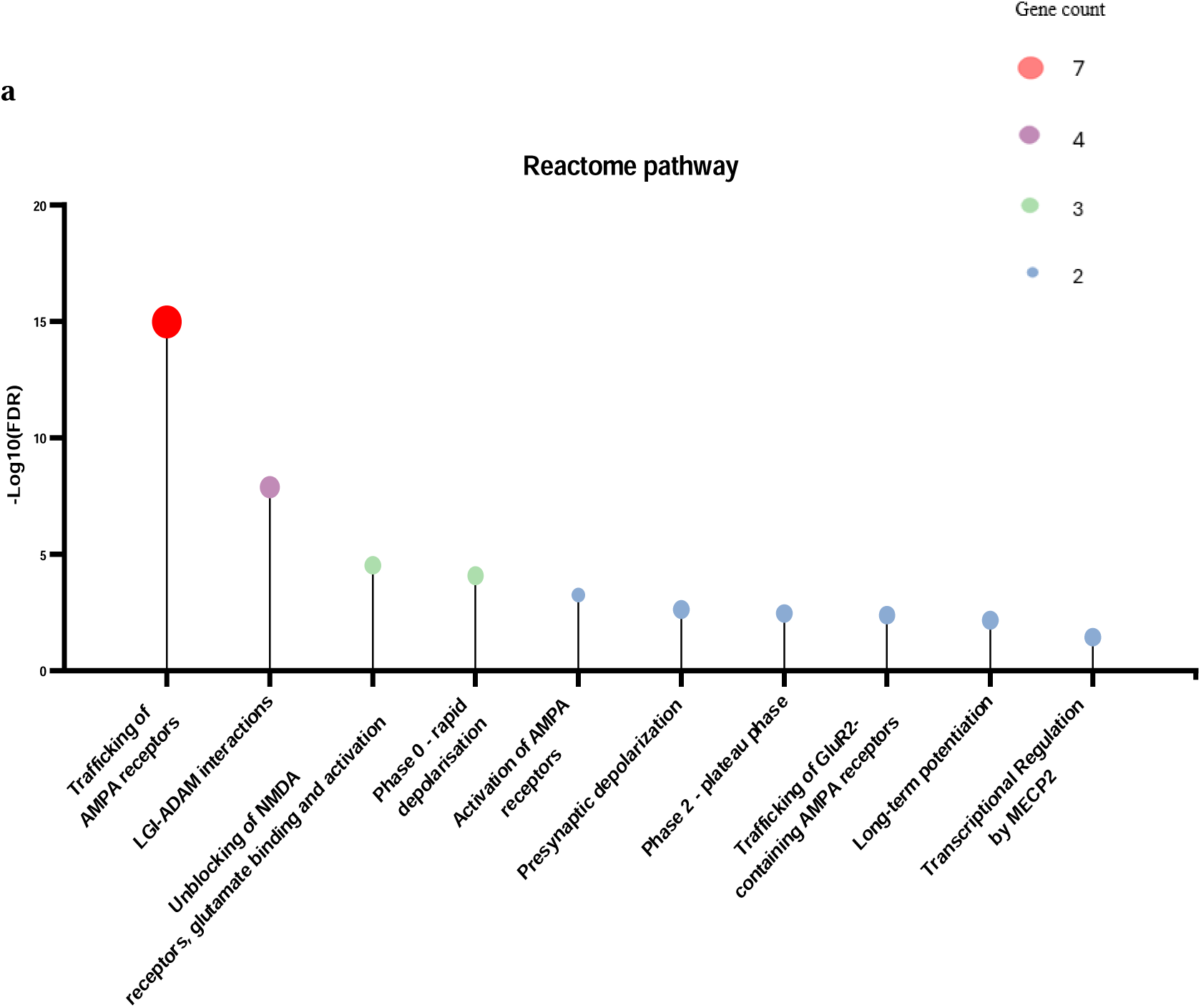
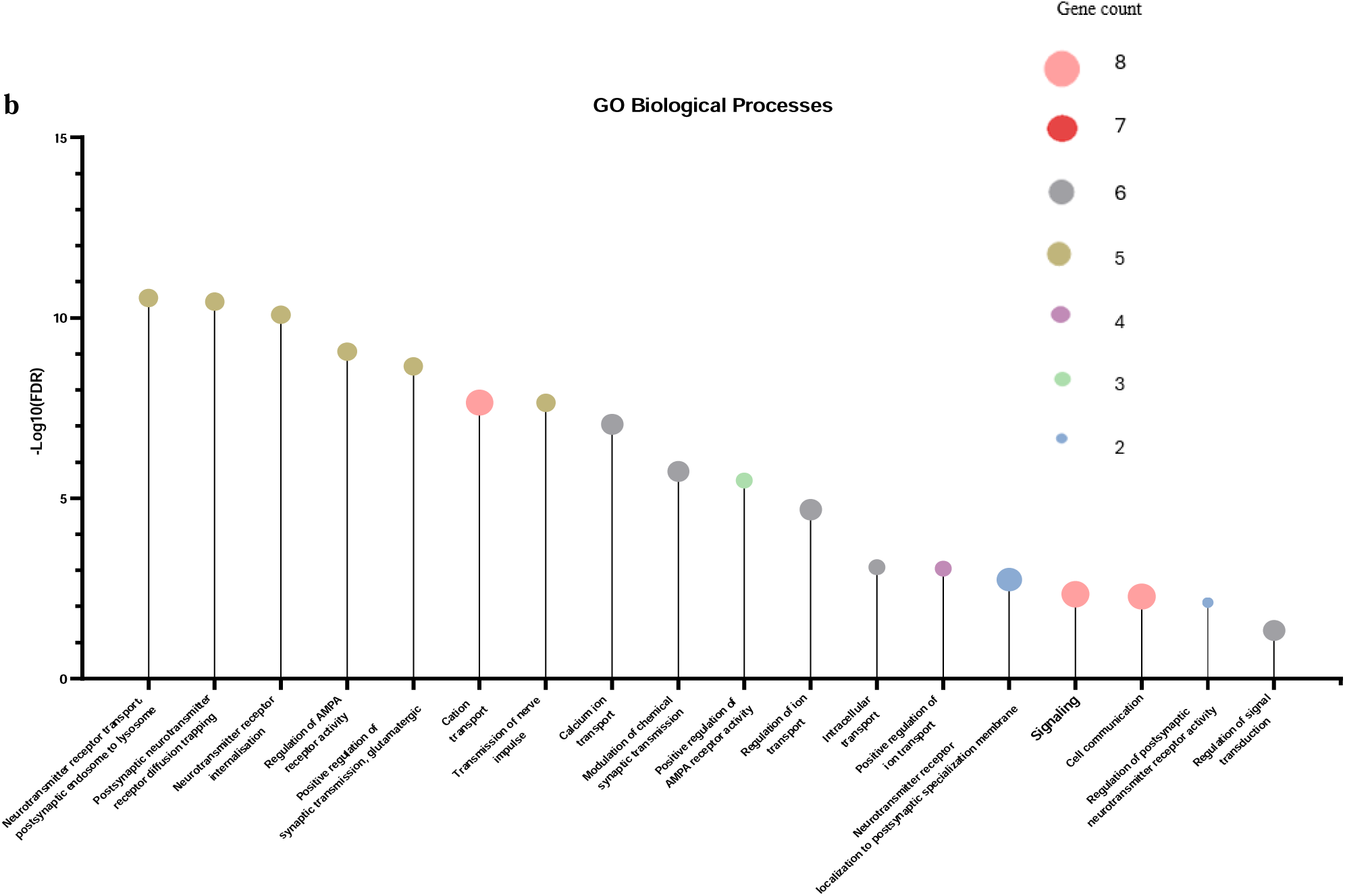
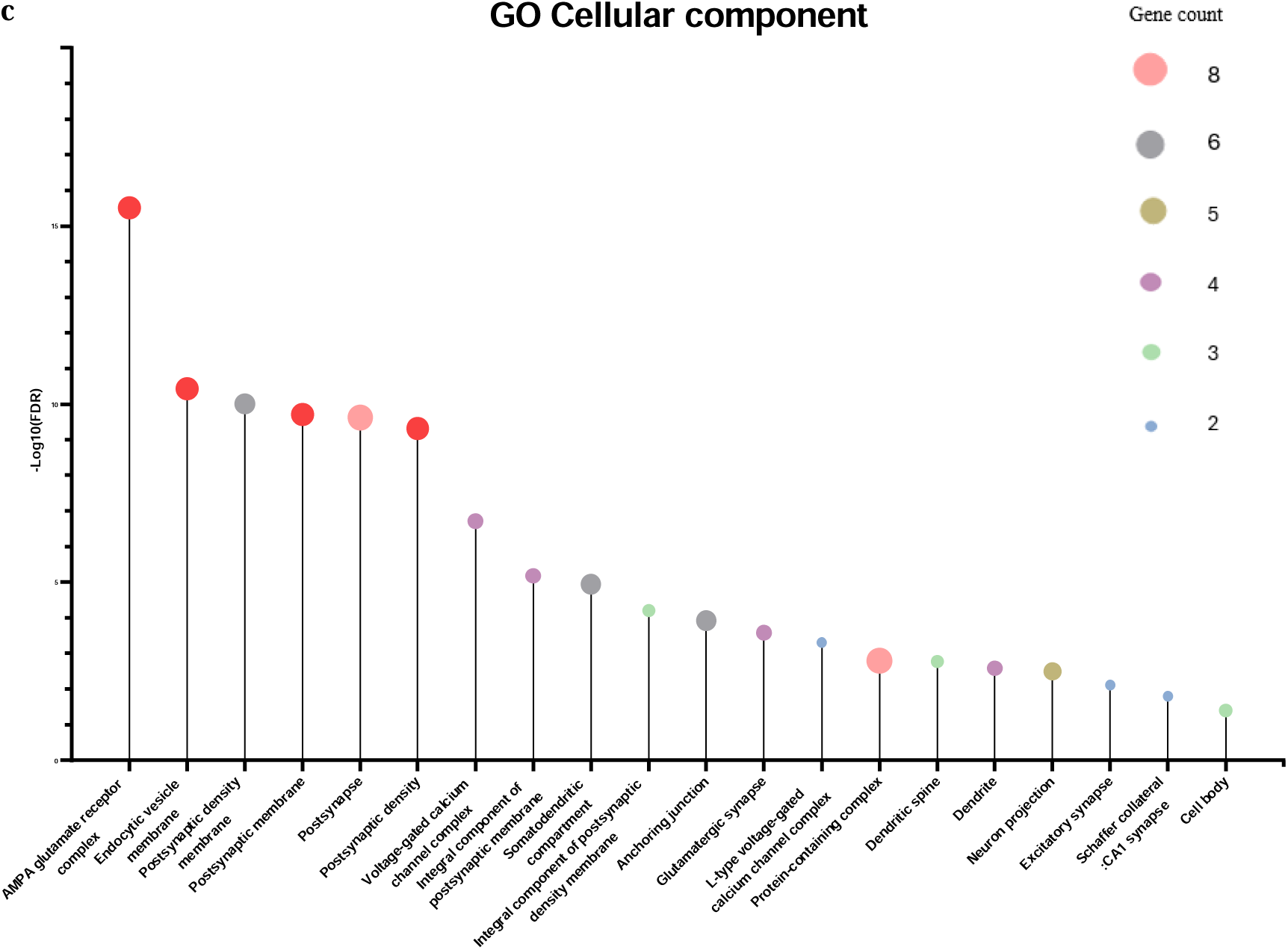
Functional enrichment analysis of the expanded hub proteins using Cytoscape and visualized using GraphPad Prism. (a) Reactome pathway enrichment: This analysis highlights a dominant overrepresentation of AMPA receptor trafficking and NMDA receptor activation, thus showing a bias towards synaptic potentiation. (b) Biological Process (Gene Ontology) enrichment: This analysis shows enrichment for both neurotransmitter receptor internalisation and the positive regulation of AMPA receptor activity, indicating an initiation of endocytosis alongside dominant receptor stabilization. (c) Cellular Component (Gene Ontology): The hub proteins are involved in the activation of the AMPA glutamate receptor complex, localisation of receptors at the postsynaptic membrane, and also in endocytic vesicle transport indicating an impaired postsynaptic receptor trafficking.

The GO profiling further validates the metaplastic mechanism lead by AMPAR trafficking. GO-BP and GO-CC confirmed the presence of ‘neurotransmitter receptor internalisation’, ‘endocytic vesicle membrane’ and ‘postsynaptic endosome to lysosome’ transport, indicating the initiation of LTD-associated receptor internalisation machinery (Fig 4b and 4c).

### 3.3 Targeted Connectome of CACNG8 and GRIA2 Identifies Auxiliary Complexes Driving AMPAR Stabilization

Having established that this core topological module is biased towards AMPA receptor trafficking and stabilization, the structural complexes that could be involved in mediating this polarity shift were identified by constructing a targeted sub-network of CACNG8 and GRIA2 as specific seed nodes (Fig 5a). These nodes were selected based on established literature stating that physical interactions between these proteins kinetically alters the AMPAR causing a delayed accumulation of current flux under continuous glutamate exposure ^16^. To isolate the minimal essential structural unit bridging these two seed nodes, their shared first-degree interactors was mechanism led by extracted. This specific filtering parameter naturally gave a tightly integrated, five-protein complex consisting of AMPAR subunits GRIA1, GRIA2, and GRIA4, TARP CACNG8, and the auxiliary trafficking protein Cornichon Homolog 2 (CNIH2) (Fig 5b). CNIH2 regulates the trafficking and gating properties AMPARs, promotes their targeting to the cell membrane and synapses, and modulates their gating properties by regulating their rates of activation, deactivation and desensitization ^17^.

**Fig 5:**
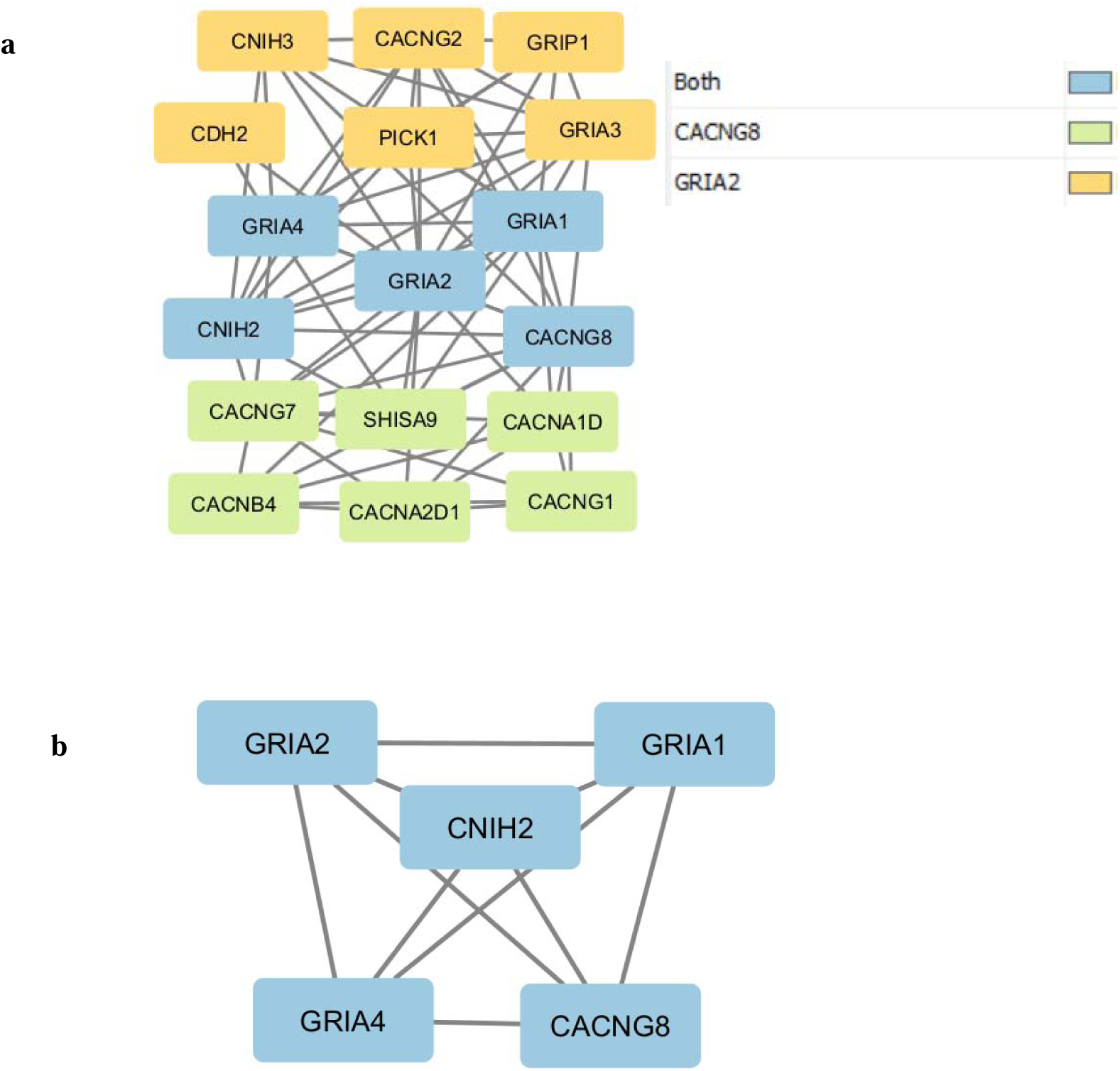
PPI network of CACNG8 and GRIA2. (a) Nodes are color-coded by their network origin (yellow: GRIA2-associated; green: CACNG8 associated; blue: shared intersecting nodes). This network has 17 nodes and 71 edges. (b) Isolated network of the intersecting nodes consisting of 5 nodes and 10 edges.

### 3.4 Topological Integration Reveals Structural Isolation of the LTD Phosphatase Machinery

Integration of calcineurin subunits, PPP3CA and PPP3CB, against the established 8-hub AMPA receptor-stabilization network (Fig 3b), resulted in a PPI network with CAMK2A positioned as a dominant central link with direct structural connections to the AMPAR subunits and auxiliary TARPs. The calcineurin subunits failed to show any direct connection to the central AMPAR complex and were connected to the network only indirectly through CAMK2A (Fig 6).

**Fig 6:**
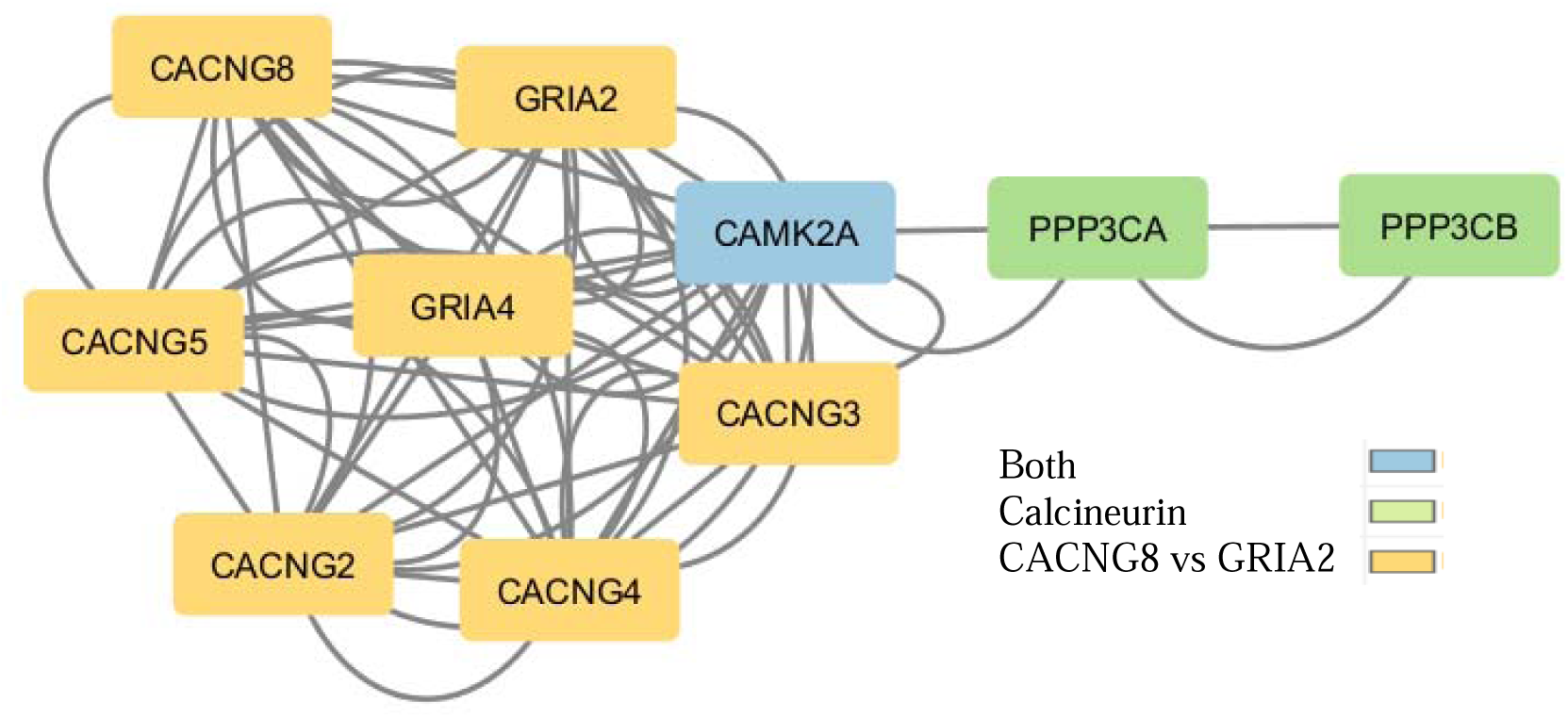
A PPI network evaluating the integration of the calcineurin catalytic subunits (PPP3CA and PPP3CB; green) into the core AMPA receptor-stabilization hub (yellow). The interactome demonstrates that the LTP-associated kinase CAMK2A (blue) is centrally embedded with dense, direct structural connections to the AMPAR subunits (GRIA2, GRIA4) and auxiliary TARPs. In contrast, the LTD-associated calcineurin subunits are indirectly connected via CAMK2A.

### 3.5 Consolidated connectome visualizes the structural basis of the metaplastic switch

To visualize the complete proposed interactome framework underlying bidirectional plasticity, a final, comprehensive PPI network was constructed (Fig 7). This unified interactome integrates all the critical proteins identified across the sequential analyses, mapping the structural relationships between the initial baseline LTD-epilepsy hubs (PLCB1, CACNA1A, GRIA2), the 8-hub core after integrating GRIN2B into the network, the targeted network (CAMK2A, GRIA4, GRIA2, TARPs and CNIH2), and the LTD-associated phosphatase machinery (PPP3CA and PPP3CB).

**Fig 7:**
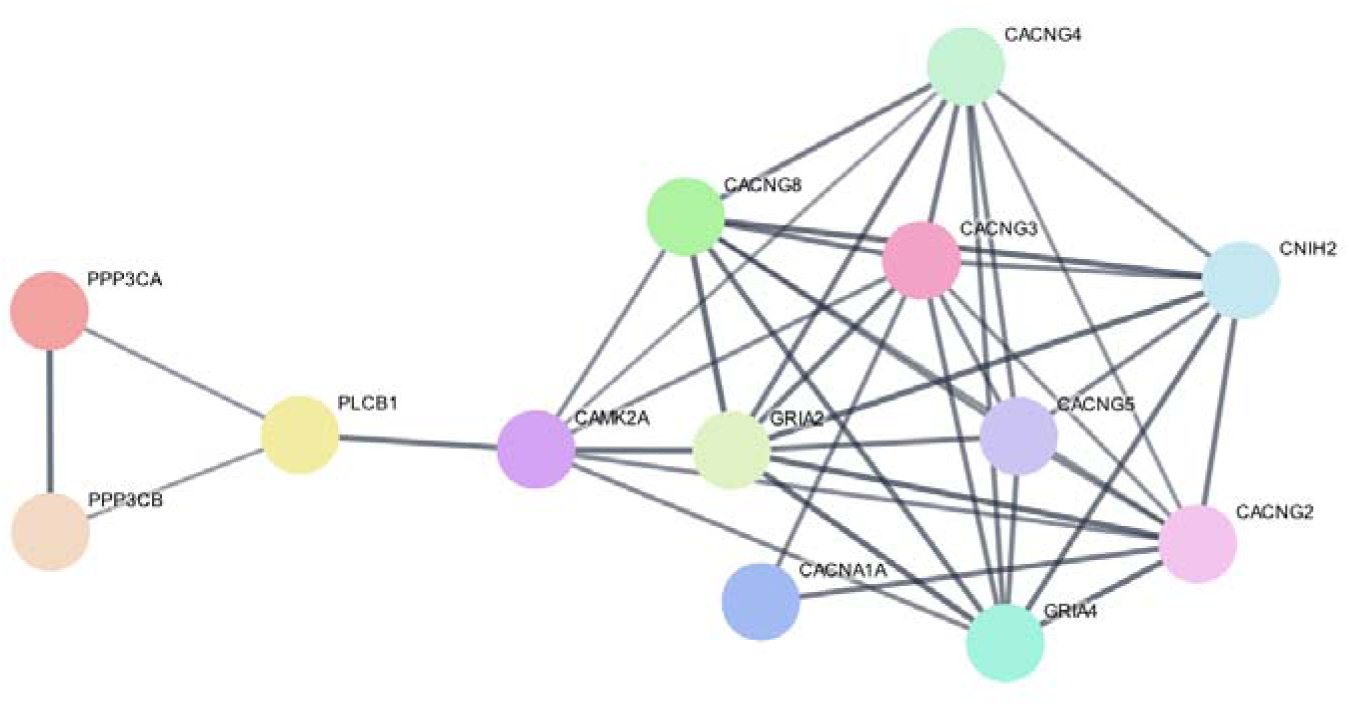
Consolidated PPI network using all functionally significant modules identified throughout the stepwise analysis. This network has 13 edges and 40 nodes.

The evaluation of this consolidated network provides a structural topography of the synapse’s vulnerability to metaplastic shifts. This interactome has two distinct functional domains, one domain dominated by AMPAR subunits and auxiliary proteins, and the other domain with LTD machinery, with CAMK2A and PLCB1 acting as the molecular bridge between these two domains.

### 3.6 Transcriptomic Cross-Validation of the Metaplasticity Hubs

To determine whether the structural dominance of the AMPA receptor-stabilizing core is reflected at the transcriptomic level in human epilepsy, the expression profiles of the identified hub genes, CNIH2, and their extended network interactors were analysed using the GSE256068 dataset.

The transcriptomic analysis revealed a significant and consistent upregulation of critical auxiliary and stabilizing proteins across both cortical and hippocampal epileptic tissues. Most notably, the auxiliary trafficking protein CNIH2, which the connectome identified as a direct structural driver of prolonged AMPAR retention, was significantly upregulated in both the cortex (log2FC = 0.765, adj.P < 0.05) and the hippocampus (log2FC = 0.776, adj.P < 0.05). Similarly, the TARP member CACNG2 (Stargazin) was also seen to be upregulated in both tissue types (Cortex log2FC = 0.621; Hippocampus log2FC = 0.593, adj.P < 0.05) (Table 1).

**Table 1:** Differential expression of core network hubs and highly dysregulated first-degree interactors in the Cortex and Hippocampus (Data from NCBI GEO: GSE256068).

| **Gene**<br>**Symbol** | **Functional Role within Network** | **Cortex (log2FC,**<br>**adj.P < 0.05)** | **Hippocampus**<br>**(log2FC,**<br>**adj.P < 0.05)** |
| --- | --- | --- | --- |
| CNIH2 | Auxiliary trafficking / AMPAR retention | 0.765 | 0.776 |
| CACNG2 | TARP / Auxiliary subunit | 0.621 | 0.593 |
| CACNG4 | TARP / Auxiliary subunit | 0.831 | n.s. |
| CAMK2A | Central LTP-associated kinase | 0.653 | n.s. |
| GRIA1 | AMPA Core Subunit | n.s. | -0.597 |
| GRIA2 | AMPA Core Subunit | n.s. | -0.726 |
| DLG4 | Postsynaptic scaffolding (First-degree interactor) | 0.624 | n.s. |
| GRIP1 | Synaptic scaffolding (First-degree interactor) | 0.69 | n.s. |
| CALML3 | Calcium signalling (First-degree interactor) | n.s. | 1.486 |

*Note: "n.s." (Not Significant) indicates that while the transcript was detected, it did not pass the defined threshold for differential expression (adj. P-value < 0.05 and absolute log2FC > 0.5) in that specific tissue.*

In the cortical tissues, the potentiation-driving kinase CAMK2A was significantly upregulated (log2FC = 0.653, adj.P < 0.05), alongside the auxiliary TARP CACNG4 (log2FC = 0.831, adj.P < 0.05). While core AMPA receptor subunits were either below the DEG threshold in the cortex or significantly downregulated in the hippocampus (*GRIA1* log2FC = −0.597; *GRIA2* log2FC = −0.726), an evaluation of their first-degree network interactors revealed significant structural dysregulation as synaptic scaffolding proteins such as DLG4 (log2FC = 0.624, adj.P < 0.05) and GRIP1 (log2FC = 0.690, adj.P < 0.05) were significantly upregulated in the cortex, and calmodulin-related signalling interactors (CALML3, log2FC = 1.486; CALML6, log2FC = 0.734, adj.P < 0.05) were highly elevated in the hippocampus.

Collectively, these transcriptomic profiles strongly validate the topological model, demonstrating that the structural auxiliary machinery responsible for capturing and stabilizing AMPA receptors (CNIH2, CACNG2, and scaffolding interactors) is pathologically overexpressed in the human epileptic brain.

## 4. Discussion

Impairment in learning and memory and cognitive functions are comorbid conditions found in patients with epilepsy and this is largely driven by the pathological alterations in synaptic functions. Electrophysiological studies in epileptic brain slices have demonstrated that epileptic synapses undergo a GluN2B-dependent metaplastic shift where LFS induces LTP instead of LTD. However, the downstream molecular rewiring driving this impairment remains unresolved. This study utilized a comprehensive *in silico* connectome approach to bridge the gap between this electrophysiological findings and synaptic remodelling to understand the molecules that could be potentially involved in this plasticity shift.

This network analysis demonstrates that the failure of LTD in the epileptic brain is not due to a failure of initiation mechanisms but rather a failure of consolidation. This is evident from the functional enrichment of the GRIN2B-anchored network wherein there were LTD-initiating functional terms such as ‘neurotransmitter receptor internalisation’ and ‘endocytic vesicle transport’^18,19^. However, if LTD had been successfully consolidated in this network, there would be GO terms associated with permanent synaptic weakening such as ‘actin depolarisation’, ‘ubiquitin-dependent receptor degradation’, and the ‘negative regulation of AMPA activity’ ^19–22^. The absence of these terms indicates that although LTD is initiated, it is functionally dominated by LTP-associated mechanisms, specifically the positive regulation of AMPA receptors and postsynaptic diffusion trapping.

The targeted sub-network using CACNG8-GRIA2 provides a deeper insight into the possible downstream molecules. The emergence of the auxiliary trafficking protein CNIH2 of the cornichon protein family, integrated alongside the TARPs and AMPARs signifies that this postsynaptic phenomenon could be due to the synaptic remodelling occurring as a result of increased surface expression of AMPARs facilitated by transport proteins. Cornichon proteins have been known to function as a powerful auxiliary subunit that actively promote surface expression of AMPAR and drastically slow their deactivation kinetics ^17,23–25^. By forming a CNIH2-TARP complex, the epileptic synapse prevents the completion of LTD by preventing the internalisation of the receptor and also establishes a physical architecture supporting prolonged receptor kinetics ^23,24^.

The consolidated network made using all the predicted nodes further illustrates an enzymatic key in the metaplasticity shift. The interactome is distinctly split into two functional domains: one containing AMPARs and TARPs, and one for the calcineurin complex, with both of these domains connected by CAMK2A and PLCB1. The structural segregation of calcineurin from the core AMPA receptor network directly could explain the inability of the synapse to consolidate LTD. In healthy synapses, for bidirectional plasticity, calcineurin must directly access and dephosphorylate AMPARs to initiate endocytic removal ^18,26,27^. However, based on the network’s topological segregation (Fig 7), the calcineurin complex lacks direct connectivity to the AMPARs. As a consequence, even if a low-frequency stimulus successfully activates calcineurin, its topological segregation from the AMPARs physically prevents the complete execution of LTD. Furthermore, the CAMK2A-PLCB1 bridge could indicate that when an LTD-inducing stimulus triggers calcium influx, this calcium must navigate through the PLCB1-CAMK2A axis to reach the calcineurin complex. But as CAMK2A is positioned directly at the gate of the receptor complex, this could be the reason why the LFS triggers CAMK2A activation and subsequent AMPAR stabilization ^28–31^.

It is important to note that PPI networks derived from text-mining and databases represent the physiological interactome blueprint rather than a disease-specific physical state. The network mapping shows that the baseline architecture of the synapse segregates calcineurin from the AMPA receptors, making the LTD process susceptible to impairment. However, the actual pathological rewiring is executed at the transcriptional level. To investigate the transcriptional environment supporting this disease-specific execution, transcriptomic analysis using human epileptic tissue (GSE256068) was evaluated. While it is well established that bulk mRNA expression is not a direct substitute for final protein abundance or subcellular localization, the RNA-sequencing data indicates that the epileptic brain pathologically exploits this interactome blueprint by actively overexpressing the specific structural components required to build the stabilization complex. The significant upregulation of the auxiliary trafficking protein CNIH2, alongside the TARP member CACNG2, across both cortical and hippocampal tissues supports the model of a persistent active synaptic remodelling that increases the excitatory current. Furthermore, the upregulation of the potentiation-driving kinase CAMK2A and essential postsynaptic scaffolding interactors (such as DLG4 and GRIP1) aligns with the topological findings. This overexpression indicates a coordinated cellular signalling aimed at producing the requisite building blocks to physically localise AMPARs to the postsynaptic density in an active conformation. Consequent proteomic studies would be necessary to support the hypothesis that the disrupted CNIH2-TARP-AMPAR complex signalling causes the impairment of LTD in epileptic synapses.

At first glance, the significant downregulation of core AMPA receptor subunits (*GRIA1*, *GRIA2*) in the hippocampus appears to contradict synaptic potentiation. However, this transcriptional reduction is highly consistent with a compensatory homeostatic response to network hyperexcitability ^32,33^. Because the pathological upregulation of CNIH2 and TARPs kinetically traps existing AMPA receptors at the postsynaptic density and the neuron experiences sustained excitatory drive. To protect against excitotoxicity, the cell attempts to scale down synaptic strength by suppressing the *de novo* global transcription of core receptor subunits ^34^. Hence, this highlights that the metaplastic shift in epilepsy is not driven by a simple overproduction of receptors, but rather by a pathological post-translational scaffolding mechanism that locks existing receptors in place, rendering the cell’s homeostatic transcriptional downregulation ineffective.

Ultimately, this network topology suggests viewing epilepsy not only as a chemical imbalance of excitatory and inhibitory neurotransmission, but also as a nanoscale structural synaptopathy. The epileptic postsynaptic density physically rewires itself, constructing a TARP/CNIH2 architecture around excitatory receptors while physically disabling the LTD machinery. This structural basis of metaplasticity helps explain the complexity of cognitive comorbidities in epilepsy.

## 5. Conclusion

This study successfully maps the downstream molecular architecture/signalling that could be responsible for the pathological synaptic plasticity shift in the epileptic synapses. By employing a stepwise *in silico* connectome approach followed by a transcriptomic analysis, this study demonstrates that the impairment of LTD is not caused by an inability to initiate endocytic signalling, but by a distinct structural rewiring that physically prevents its consolidation. The epileptic postsynaptic interactome was topologically polarized: the essential LTD machinery (calcineurin) was functionally uncoupled from the receptor core, while the potentiation-driving kinase (CAMK2A) maintains dominant central connectivity. Concurrently, the integration of auxiliary TARPs and CNIH2 creates a stabilization complex that kinetically traps AMPA receptors at the membrane, functionally overriding LTD machinery and forcing the synapse into continuous potentiation. Ultimately, these findings translate electrophysiological phenotypes into a structural framework, identifying CNIH2-TARP-AMPAR complex as highly specific, novel molecular targets for therapeutic intervention aimed at rescuing cognitive comorbidities in epilepsy.

